# Functional Recovery of the Germ Line Following Splicing Collapse

**DOI:** 10.1101/2021.06.04.447173

**Authors:** Wei Cao, Christopher Tran, Stuart K. Archer, Sandeep Gopal, Roger Pocock

## Abstract

Splicing introns from precursor-messenger RNA (pre-mRNA) transcripts is essential for translating functional proteins. Here, we report that the previously uncharacterized *Caenorhabditis elegans* protein MOG-7, acts as a pre-mRNA splicing factor. Depleting MOG-7 from the *C. elegans* germ line causes intron retention in the majority of germline-expressed genes, impeding the germ cell cycle, and causing defects in nuclear morphology, germ cell identity and sterility. Despite the deleterious consequences caused by MOG-7 loss, the adult germ line can functionally recover to produce viable and fertile progeny when MOG-7 is restored. Germline recovery is dependent on a burst of apoptosis that likely clears defective germ cells, and viable gametes generated from the proliferation of germ cells in the progenitor zone. Together, these findings reveal that MOG-7 is essential for germ cell development, and that the germ line is able to functionally recover after a collapse in RNA splicing.

## INTRODUCTION

In many species, gametes are produced throughout reproductive life from adult stem cells housed in the germ line. Correct developmental control of germ cell fates is crucial for generating viable gametes and species propagation. As such, mechanisms that control germ cell proliferation, differentiation, survival and sex determination are directed by highly regulated networks of gene expression.

The *Caenorhabditis elegans* hermaphrodite germ line consists of a pair of U-shaped syncytial tubes organized in a distal-proximal fashion. A population of germline stem cells (GSCs) and their proliferating progeny are located at the distal end of each tube, with mature gametes at their proximal ends. The GSCs reside in a specialized niche supported by Notch signals from somatic cells ^1, 2^. GSCs within the niche both self-renew to maintain the stem cell population and produce germ cells that exit the niche and begin the differentiation process. Further proximally, germ cells proceed through meiotic pachytene prior to gametogenesis where they transiently generate sperm from the L3 larval stage, before permanently switching to oogenesis around the L4/adult transition ^3^. During the late pachytene phase of the oogenic program, approximately half the germ cells are removed by physiological apoptosis ^4^. These apoptosed germ cells are thought to function as nurse cells by providing cytoplasmic components to maturing oocytes ^4^. Germ cell apoptosis is also triggered by DNA damage, environmental stress and pathogen exposure ^5–7^.

Post-transcriptional gene regulation is critically important for controlling germ cell fates in the *C. elegans* germ line. In particular, precursor-messenger RNA (pre-mRNA) splicing has a strong influence on the transition between germ cell proliferation/differentiation and the sperm-oocyte switch ^8, 9^. Splicing of intervening non-coding sequences (introns) from protein-encoding sequences of eukaryotic genes is a highly conserved and complex process (reviewed in ^10^). The splicing reaction occurs in a stepwise fashion within a dynamic ribonucleoprotein complex called the spliceosome, containing over 300 proteins in humans ^10^. Intron excision occurs in two sequential transesterification reactions that result in the excision of the intron lariat, and the ligation of the 3’ and 5’ exons. The spliceosome is then disassembled, and the lariat released from the intron lariat spliceosome complex (ILS), freeing the spliced mRNA for protein synthesis and enabling the spliceosome complex components and intron nucleotides to be recycled. Spliceosome disassembly is essential for efficient splicing because the spliceosome assembles *de novo* at each new round of pre-mRNA splicing ^11^. Splicing factors, encoded by *mog* genes (*m*asculinization *o*f the *g*erm line), promote *C. elegans* oogenesis: their depletion causes a failure in the sperm-oocyte switch and the generation of a germ line replete with sperm (spermatogenic germ line) ^12, 13^. The switch from spermatogenesis to oogenesis is controlled by many RNA regulatory proteins that comprise the germ line sex determination pathway ^14^. It is thought that inefficient splicing of these sex determining genes caused by mutations in *mog* genes inhibits appropriate switching of sexual fate in the germ line ^8^.

Here, we determined the function of the previously uncharacterized gene *mog-7* in the *C. elegans* germline. The *mog-7* locus encodes two protein isoforms: a long isoform containing a *Saccharomyces cerevisiae* Ntr2 (Nineteen Complex-Related Protein 2) domain and a human GCFC (GC-rich sequence DNA-binding factor) domain, and a short isoform containing only the GCFC domain. Ntr2 and GCFC domains are found in factors that regulate the disassembly of the ILS by the DEAH family ATPase/helicase Prp43 ^15, 16^. We found that MOG-7 is expressed throughout the germ line and that germline-specific removal of MOG-7 inhibits germline development and causes complete sterility; likely due to abnormalities observed in nuclear morphology and the germ cell cycle. We found that MOG-7 interacts *in vivo* with multiple splicing factors required for spliceosome disassembly. Moreover, germline-specific removal of MOG-7 causes intron retention in over 5000 germline-expressed genes. Despite the severe defects in RNA splicing and germ cell behavior, the germ line can recover from MOG-7 depletion to produce viable and fertile progeny if MOG-7 expression is restored. We observed a burst of apoptotic death in the germ line within hours of restoring MOG-7 expression, and recovery of germline function is dependent on the apoptotic caspase CED-3. Therefore, removal of defective germ cells is required for germline recovery following restoration of MOG-7 expression. In addition, germline recovery from splicing collapse requires proliferation, likely from a cohort of arrested distal germ cells. Together, we demonstrate that the *C. elegans* germ line exhibits a capacity to functionally recover following a catastrophic breakdown of RNA splicing.

## RESULTS

### F43G9.12/MOG-7 is essential for *C. elegans* germline development

In an ongoing RNA-mediated interference (RNAi) screen for nucleic acid binding factors that control germline development, we identified the uncharacterized gene *F43G9.12/mog-7*. The *mog-7* locus generates two protein isoforms - MOG-7a and MOG-7b (Fig. 1A). MOG-7a contains Ntr2 and GCFC domains, with MOG-7b housing the GCFC domain alone (Fig. 1A). Proteins containing these domains have been shown to independently control splicing in *Saccharomyces cerevisiae* (Ntr2) and humans (GCFC - also known as C2orf3) ^15, 17, 18^.

**Figure 1.**
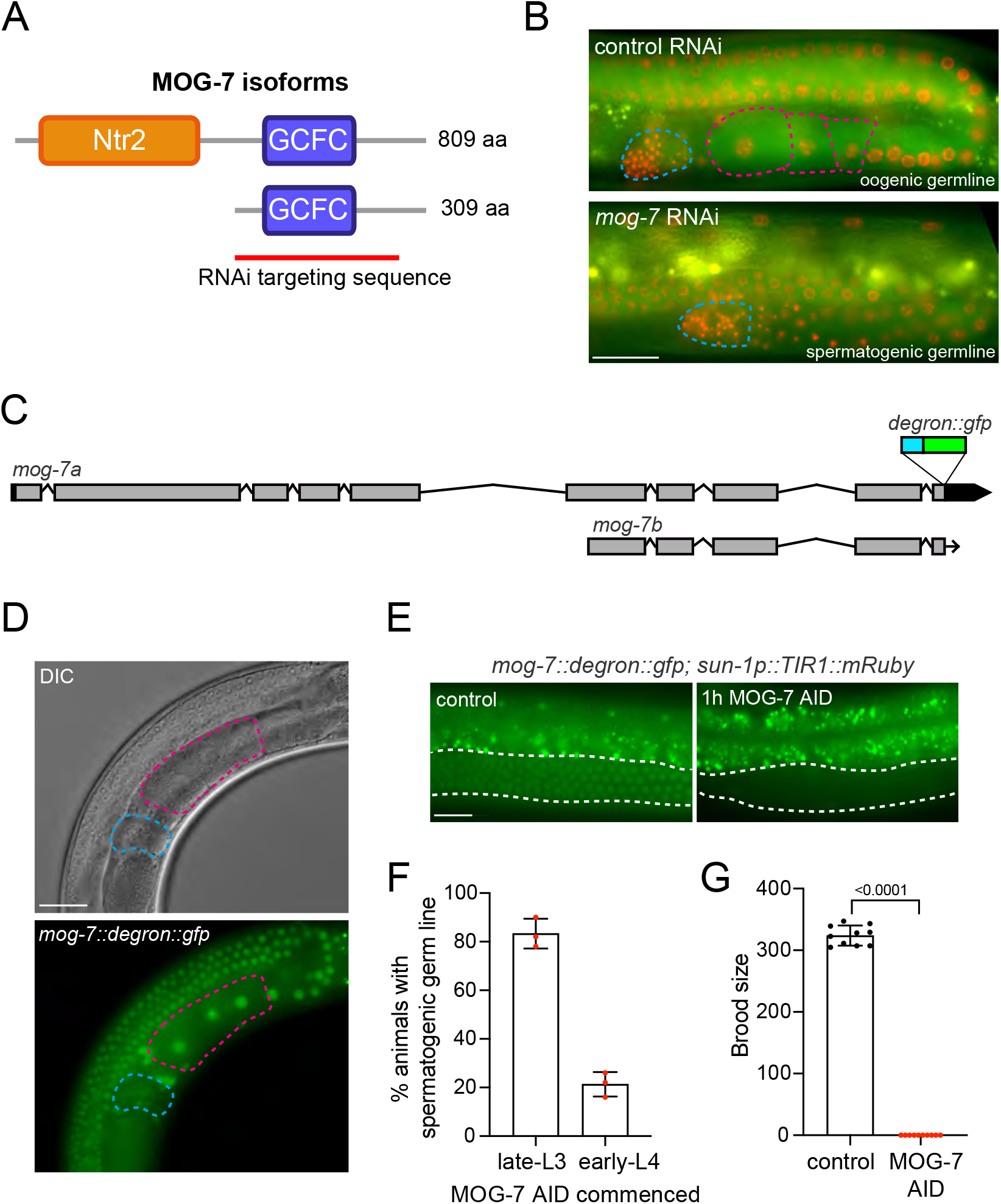
MOG-7 controls germ cell fate and fertility. (A) Protein domain of MOG-7a and MOG-7b. Ntr2 = Nineteen complex-related protein 2, pre-mRNA splicing factor domain. GCFC = GC-rich sequence DNA-binding factor 2 domain. Red line = targeting sequence in RNAi experiments. (B) Fluorescence micrographs of one-day adult germ lines +/- *mog-7* RNAi from the L1 larval stage (control = L4440 vector). Micrographs show chromosomes in red (mCherry::HIS-58) and plasma membrane in green (GFP::PH(PLC1delta1)) ^45^. Spermatheca = blue dashes. Oocytes = pink dashes. Scale bar = 20 μm. (C) *mog-7* genomic structure showing the insertion site of the *degron::gfp* coding sequence at the C-terminus of both isoforms. (D) Differential interference contrast (DIC) and fluorescence micrographs of *mog-7::degron::gfp* expression in an adult hermaphrodite. Spermatheca = blue dashes. Oocytes = pink dashes. Scale bar = 20 μm. (E) Fluorescence micrographs of *mog-7::degron::gfp* expression +/- auxin treatment for 1h (ethanol used as a control). *sun-1p::TIR1::mRuby* = germline-specific auxin-inducible degron strain. Dashed line = germ line region. Scale bar = 10 μm (F) Percentage of spermatogenic adult hermaphrodite germ lines of *mog-7::degron::gfp; sun-1p::TIR1::mRuby* exposed to auxin from the late-L3 or late-L4 stage. Data expressed as mean ± s.e.m. and statistical significance was assessed by Welch’s t-test. n = 40 per replicate. (G) Quantification of brood size of *mog-7::degron::gfp; sun-1p::TIR1::mRuby* hermaphrodites exposed to ethanol or auxin from the early L4 stage. Data expressed as mean ± s.e.m. and statistical significance was assessed by Welch’s t-test. n = 10.

We explored the function of *mog-7* in the germ line by RNAi targeting of both isoforms (Fig. 1A-B and Fig. S1). *mog-7* RNAi knockdown from the L1 larval stage induced a spermatogenic germ line in approximately 35% of P0 animals (Fig. 1B and Fig. S1A). This phenotype corresponds with other studies of splicing factors in the *C. elegans* germ line that also control the sperm-oocyte switch ^8, 12, 13, 19–21^. *mog-7* L1 RNAi also resulted in low broods with most F1 progeny dying during embryogenesis (Fig. S1B). *mog-7* RNAi from the L4 larval stage enabled hermaphrodites to generate more progeny than L1 knockdown however few embryos hatched into larvae (Fig. S1C). Thus, *mog-7* is critical for fertility and embryonic viability.

### MOG-7 autonomously controls germline development

To examine the endogenous expression pattern of MOG-7, we used CRISPR-Cas9 to knock-in a *degron::green fluorescent protein* (*gfp*) coding sequence at the 3’ end of the *mog-7* gene, thereby tagging both MOG-7 isoforms (Fig. 1C) ^22^. We inserted the degron sequence to enable acute removal of endogenous MOG-7 using the auxin-inducible degron (AID) system ^23^. MOG-7::degron::GFP expression was detected in germline nuclei throughout development, commencing in the primordial germ cells of L1 larvae through to oocytes and sperm in adults (Fig. 1D-E and Fig. S2). Somatic expression of MOG-7::degron::GFP was detected in embryos from the 2-cell stage and throughout worm development in most, if not all, cells (Fig. S2). MOG-7::degron::GFP expression was also detected by Western blot confirming the presence of both MOG-7 isoforms (Fig. S2F). We found that insertion of the degron::GFP sequence had no overt effect on MOG-7 germline function as shown by germ cell counting and brood size analysis (Fig. S3). In addition, incubating animals on auxin-containing media did not affect germ cell number (Fig. S3).

To validate the efficacy of the MOG-7::degron::GFP allele for functional studies, we introduced a transgene that expresses the *Arabidopsis thaliana* F-box protein called transport inhibitor response 1 (TIR1) in the germ line (*sun-1p::TIR1::mRuby*) of *mog-7::degron::gfp* animals ^23^. In the presence of auxin, TIR1 targets AID-tagged proteins for ubiquitin-dependent proteasomal degradation ^23^. The *sun-1p::TIR1::mRuby* transgene had no detectable effect on germline function either alone or in the presence of *mog-7::degron::gfp* (Fig. S3). We found that exposing *mog-7::degron::gfp; sun-1p::TIR1::mRuby* adult hermaphrodites to auxin for 1h depleted MOG-7::degron::GFP specifically in the germ line, without detectable changes in the soma (Fig. 1E and Fig. S3E). Therefore, MOG-7::degron::GFP can be rapidly removed using the AID system.

Our initial RNAi analysis showed that *mog-7* knockdown causes defects in germ cell fate and fecundity (Fig. 1B and Fig. S1). However, this phenotype may be caused by a non-autonomous function for *mog-7* from somatic tissue. As we observed germline-specific loss of MOG-7::degron::GFP expression in *mog-7::degron::gfp; sun-1p::TIR1::mRuby* animals following auxin exposure (called MOG-7 AID from here) (Fig. 1E and Fig S3E), we used this tool to examine the autonomous role of *mog-7* in the germ line. *mog-7* RNAi from the L1 larval stage caused a spermatogenic germ line in ~35% of animals (Fig. 1B and Fig. S1A). As spermatogenesis begins in late L3 larvae ^24^, we performed continuous MOG-7 AID from this stage and found that ~80% of animals had spermatogenic germ lines in the first day of adulthood (Fig. 1F). In contrast, MOG-7 AID from the early L4 stage (6h after late L3) resulted in mostly oogenic germ lines (Fig. 1F). This reveals the importance of MOG-7 in determining the sexual fate of germ cells before the L4 stage. Next, we determined the effect of MOG-7 AID on fecundity after the sperm-oocyte switch with continuous auxin treatment from the early L4 stage. We found that all MOG-7 AID animals were sterile, indicating a severe defect in germline health (Fig. 1G). The germ cell fate and fecundity phenotypes caused by MOG-7 AID were stronger than in our RNAi experiments, likely because RNAi does not completely remove *mog-7* mRNA. Taken together, these data show that MOG-7 is a critical, autonomous regulator of germline development.

### MOG-7 is acutely required to maintain germ cell health

We next explored the cellular basis of germline sterility caused by MOG-7 AID. To dissect the function of MOG-7 in regulating germ cell behavior, we grew hermaphrodites to 1-day adults and then performed MOG-7 AID for 6h. We then extracted germ lines and examined germ cell nuclear morphology by staining DNA with 4’,6-diamidino-2-phenylindole (DAPI) (Fig. 2A-C). In the progenitor zone (PZ), we observed germ cells with bright florets of DAPI-stained DNA, some of which with large nuclei, suggesting cell cycle arrest (Fig. 2A) ^25^. The transition zone is normally dominated by nuclei in early meiotic prophase, which have a crescent-shaped chromosome morphology indicative of meiotic chromosome pairing ^26^. We found that MOG-7 AID caused severe defects in germ cell morphology such that few crescent-shaped nuclei were recognizable, suggesting defects in chromosome packaging (Fig. 2B). As germ cells advance into the pachytene region, synapsed homologous chromosomes form compact ribbons (Fig. 2C). After MOG-7 depletion, pachytene nuclei are spatially disorganized and have irregular morphology (Fig. 2C). Together, these data reveal that depletion of *mog-7* causes defects in germ cell nuclear morphology throughout the germ line.

**Figure 2.**
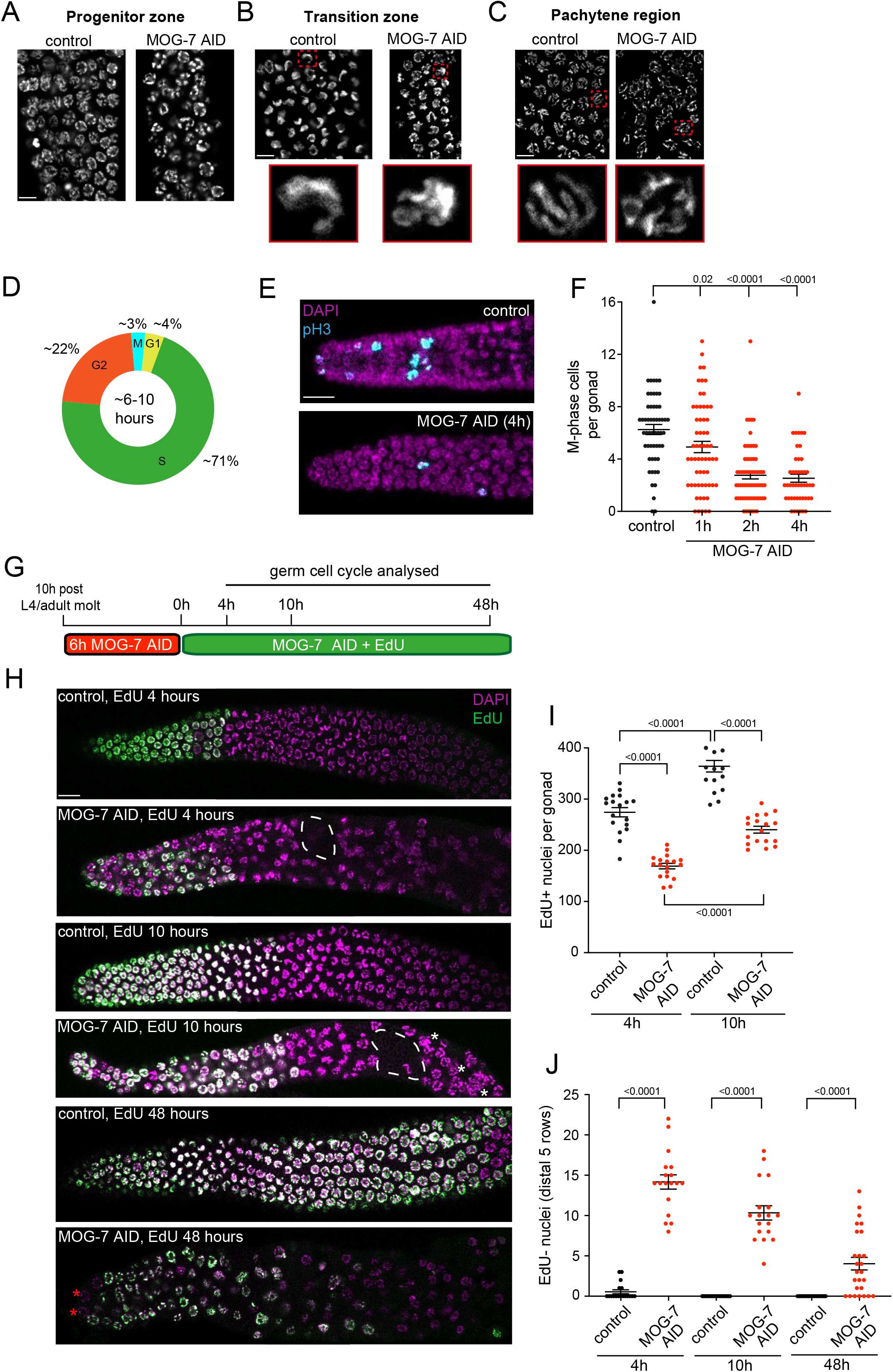
MOG-7 depletion causes defects in nuclear morphology and germ cell behavior. (A-C) DAPI-stained germ cell nuclei in the progenitor zone (A), transition zone (B), pachytene region (C) of *mog-7::degron::gfp; sun-1p::TIR1::mRuby* 1-day adult hermaphrodites exposed to ethanol or auxin for 6h. Hatched red boxes highlight individual nuclei magnified below where appropriate. Scale bars = 5 μm. (D) Schematic of germ cell cycle length and percentage of germ cells at each cell cycle stage in an adult hermaphrodite. (Taken from ^39^). (E-F) Fluorescence micrographs (E) and quantification (F) of M-phase cells in distal germ lines of *mog-7::degron::gfp; sun-1p::TIR1::mRuby* 1-day adult hermaphrodites exposed to ethanol or auxin for 1h, 2h and 4h. Germ lines stained with DAPI to visualize DNA (magenta) and anti-phospho-histone H3 (pH3) to visualize M-phase chromosomes (cyan). Data expressed as mean ± s.e.m. and statistical significance was assessed by one-way ANOVA multiple comparison with Sidak correction. n = 45. Scale bar = 10 μm. (G) Experimental outline for S-phase labelling of adult germ cells following *mog-7* depletion. 10h after the L4/adult molt, *mog-7::degron::gfp; sun-1p::TIR1::mRuby* hermaphrodites were continuously incubated on auxin or ethanol plates. After 6h of MOG-7 AID, animals were labelled with 5-ethynyl-2’-deoxyuridine (EdU) by feeding for 4h, 10h and 48h, and germ lines extruded and analyzed. Control animals were exposed to ethanol for the same periods. (H) Fluorescence micrographs of *mog-7::degron::gfp; sun-1p::TIR1::mRuby* 1-day adult control and MOG-7 AID hermaphrodites labelled with EdU for 4h, 10h and 48h. Germ lines stained with DAPI to visualize DNA (magenta) and labelled with EdU to detect replicated DNA (green). Areas devoid of germ cells = white dashed area. Nuclei clusters = white asterisks. EdU-negative (EdU^-^) nuclei = red asterisks. Scale bar = 10 μm. (I) Quantification of EdU^+^ nuclei in *mog-7::degron::gfp; sun-1p::TIR1::mRuby* 1-day adult hermaphrodites exposed to ethanol or auxin for 4h and 10h. Data expressed as mean ± s.e.m. and statistical significance was assessed by one-way ANOVA multiple comparison with Sidak correction. n = 17-18. (J) Quantification of EdU^-^ nuclei in *mog-7::degron::gfp; sun-1p::TIR1::mRuby* 1-day adult hermaphrodites exposed to ethanol or auxin for 4h, 10h and 48h. Data expressed as mean ± s.e.m. and statistical significance was assessed by one-way ANOVA multiple comparison with Sidak correction. n = 17-18.

We next examined whether MOG-7 depletion affects germ cell behavior by examining the cell cycle (Fig. 2D-J). To assess the proliferative state of PZ cells after MOG-7 AID, we detected cells in M-phase by staining germ lines with an antibody against phospho-histone H3 (pH3) (Fig. 2E-F). We found that the number of M-phase cells in adult germ lines was significantly reduced compared to control after 1h of MOG-7 AID (Fig. 2F). After 2h of MOG-7 AID, approximately half the number of control pH3-positive nuclei were detected and this level remained unchanged after 4h, suggesting reduced/slowing in proliferation (Fig. 2F).

To provide more granularity to potential cell cycle alterations caused by MOG-7 AID, we labelled S-phase germ cells in the PZ with the thymidine analog 5-ethynyl-2’-deoxyuridine (EdU) in a time-course experiment (Fig. 2G). To examine the ability of MOG-7-depleted germ cells to enter S-phase, we exposed 1-day adults (10h after the L4/adult molt) to MOG-7 AID for 6h, and then supplied EdU for 4, 10 and 48h while maintaining MOG-7 depletion (Fig. 2G-J). After 4h and 10h of EdU treatment, we observed fewer EdU^+^ cells in MOG-7 AID germ lines compared to control animals, suggesting a slowing of the cell cycle (Fig. 2H-I). However, there was a significant increase in EdU^+^ cells between 4h and 10h in MOG-7 AID germ lines indicating that germ cells can still enter S-phase (Fig. 2I). We also observed areas of the germ line that were devoid of nuclei and the presence of nuclei clusters, revealing that the organization of germ cells through the germ line is perturbed (Fig. 2H). We next examined S-phase entry of PZ germ cells in the 5 most distal rows that house the GSCs (Fig. 2J) ^27^. In control animals, all distal germ cells are labelled with EdU within 10h (most are in fact labelled within 4h) (Fig. 2J). However, in MOG-7 AID animals 10-15 germ cells are EdU^-^ after 10h of EdU labelling (Fig. 2J). Even after 48h of EdU incubation some germ cells were not labelled with EdU (Fig. 2H and J). Thus, germ cells in the PZ respond to MOG-7 depletion by slowing the cell cycle, with a cohort of germ cells arresting prior to S-phase. Taken together, these data reveal that MOG-7 AID causes diverse defects in germ cell behavior and morphology throughout the germ line that likely account for the sterility phenotype.

### MOG-7 interacts with multiple pre-mRNA splicing factors

To explore how MOG-7 controls germ cell behavior, we identified MOG-7::degron::GFP *in vivo* interacting proteins by immunoprecipitation (IP) and mass spectrometry (MS) (Fig. 3A). We identified five candidate proteins that interact with MOG-7::degron::GFP in each of three independent MS experiments (Fig. 3B and Table S1). Intriguingly, 4/5 candidate MOG-7 interactors are found in the ILS complex in *S. cerevisiae* (EFTU-2/Snu114p, PRP-8/Prp8p, STIP-1/Ntr1p and SKP-1/Prp45p), and one is uncharacterized (Y94H6A.3) (Fig. 3B) ^28^. To confirm the interactions identified by MS analysis, we performed independent coimmunoprecipitation experiments where V5-tagged MOG-7a or MOG-7b were coexpressed in COS-7 cells with each candidate interacting protein fused to a FLAG tag (Fig. 3C, Fig. S4 and Table S1). We confirmed that FLAG-tagged EFTU-2, PRP-8, STIP-1 and SKP-1 interact with both MOG-7 isoforms, whereas an interaction was only detected between Y94H6A.3 and MOG-7a-V5 (Fig. 3C and Table S1). One of the MOG-7 interactors we identified called SKP-1 is a component of the PRP19 NTC-related (NTR) complex that is essential for catalyzing spliceosome disassembly (Fig. 3B-C) ^29^. We found that both MOG-7 isoforms also interact with PRP-19 in mammalian cells suggesting a function for MOG-7 in controlling spliceosome disassembly (Fig. 3C and Table S1). In *S. cerevisiae*, the Ntr2 protein interacts with Ntr1 (STIP-1 in *C. elegans*) within the ILS ^28^. We therefore wondered whether the Ntr2 domain of MOG-7a can independently interact with the ILS components we detected by mass spectrometry. We therefore removed the GCFC domain from MOG-7a and co-expressed the Ntr2 domain with the ILS complex proteins (Fig. 3D and Fig. S4). We found that the Ntr2 domain of MOG-7a can interact with these proteins (Fig. 3D and Fig. S4). Together, these data reveal that both the Ntr2 and GCFC domains of MOG-7 interact with multiple proteins involved in spliceosome disassembly, suggesting a function for MOG-7 in this process.

**Figure 3.**
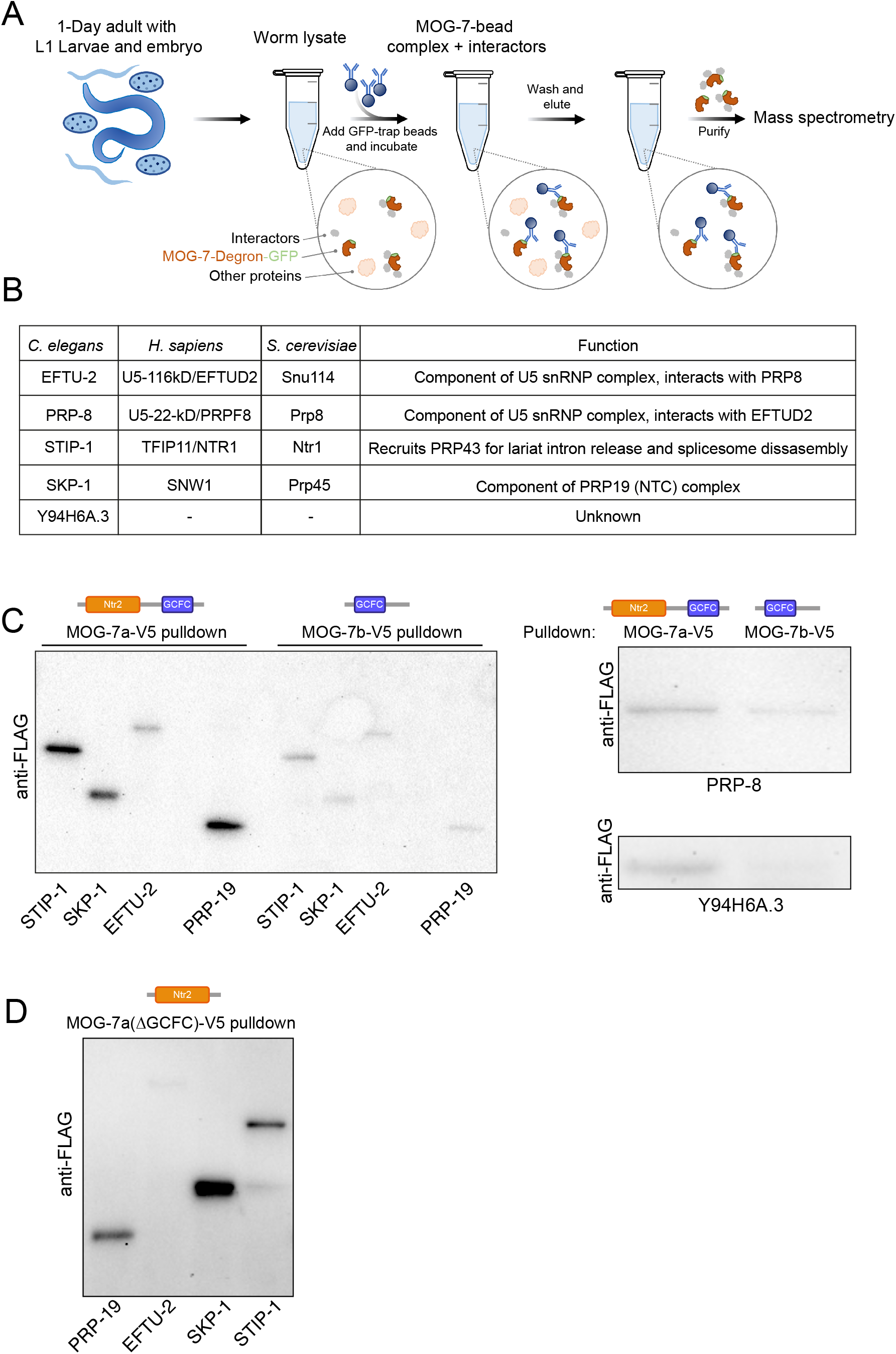
MOG-7 interacts with specific spliceosome proteins. (A) MOG-7::degron::GFP was immunoprecipitated from adult hermaphrodites, L1 larvae and embryos using magnetic beads coated with GFP-Trap. Samples were washed to remove proteins not interacting with MOG-7::degron::GFP and then analyzed by mass spectrometry after trypsin digestion. Three independent experiments were performed. (B) Proteins that reproducibly associated with MOG-7::degron::GFP in mass spectrometry experiments were core spliceosome components (EFTU-2, PRP-8, SKP-1), a spliceosome disassembly component (STIP-1) and an uncharacterized protein (Y94H6A.3). Orthologs (*H. sapiens* and *S. cerevisiae*) and known functions of these proteins are shown. (C-D) Immunoprecipitation of MOG-7a and MOG-7b (C) or MOG-7a(ΔGCFC) (D) followed by Western blot in COS-7 cells confirming the interaction of both MOG-7 isoforms with spliceosome proteins identified by mass spectrometry (n = >2 independent experiments). Predicted proteins sizes: MOG-7a-V5 (94kD), MOG-7a-V5 (36kD), FLAG-STIP-1 (94kD), FLAG-PRP-8 (272kD), FLAG-EFTU-2 (110kD), FLAG-Y94H6.A3 (21kD), FLAG-SKP-1 (60kD), FLAG-PRP-19 (53kD). Detailed analysis of interactions in Table S1.

### Acute loss of MOG-7 causes intron retention

MOG-7a contains Ntr2 and GCFC domains that regulate spliceosome disassembly in *S. cerevisiae* and humans, respectively ^15, 18^. In addition, we have shown that both MOG-7 isoforms interact *in vivo* with multiple proteins that control lariat intron release and spliceosome disassembly (Fig. 3). As recycling of these spliceosome components is a pre-requisite for each new round of splicing ^11^, we hypothesized that MOG-7 is important for efficient splicing in the *C. elegans* germline. To examine the impact of acute MOG-7 AID depletion on the transcriptome, we performed germline-specific MOG-7 AID on synchronized young adult hermaphrodites (prior to formation of embryos) for 1 or 2h followed by whole-animal RNA sequencing (Fig. 4A - see methods for detailed protocol). Ethanol treatment for 2h was used as a control. We have shown that within 1h of AID, MOG-7::degron::GFP protein is undetectable and germline phenotypes are detected (Figs. 1 and 2). Therefore, we predicted that the potential detrimental effect of MOG-7::degron::GFP loss on the transcriptome could be detected within 1h of MOG-7 depletion. Six replicates of each treatment were collected for RNA extraction and sequencing. Multi-dimensional scaling (MDS) analysis shows that each treatment group clusters together and independently of each other (Fig. S5A).

**Figure 4.**
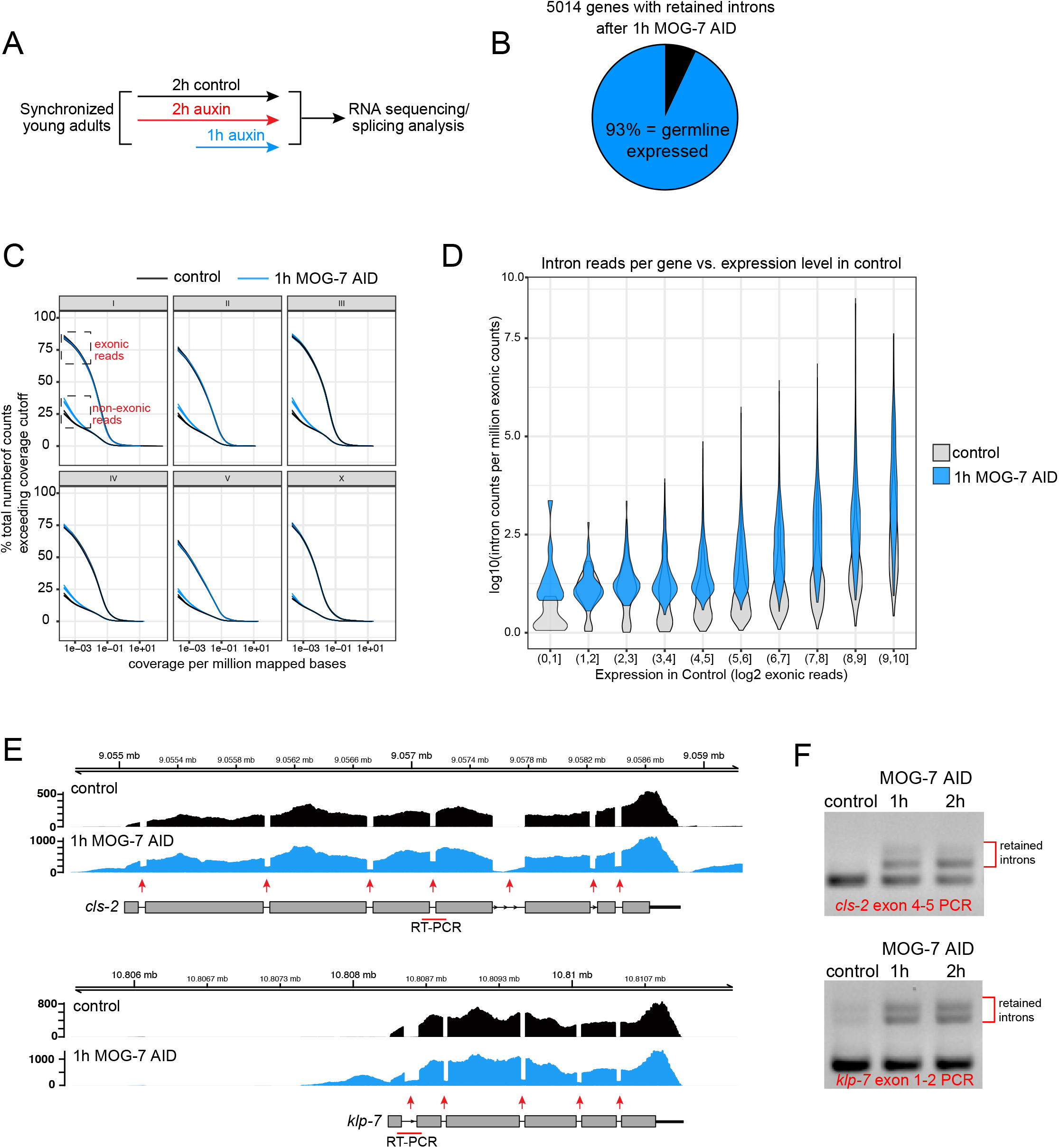
Acute loss of MOG-7 disrupts RNA splicing. (A) Experimental summary of *mog-7* depletion for RNA sequencing and splicing analysis. A population of synchronized young adult *mog-7::degron::gfp; sun-1p::TIR1::mRuby* hermaphrodites (with no embryos) were divided into three groups. Two groups were incubated either on ethanol (control) or 1 mM auxin for 2h. After 1h, the remaining group was incubated on 1 mM auxin for 1h. Six replicates of each sample were prepared for RNA sequencing and splicing analysis. (B) Summary of the number of protein-coding genes with retained introns after germline-specific MOG-7 AID (1h auxin treatment) compared to the ethanol-treated control. Intron-overlapping RNA-seq reads were aggregated at the gene level followed by analysis using EdgeR QL (Methods) with an FDR cutoff of 0.001. (C) Percentage of exonic and non-exonic base-positions (vertical axis) exceeding any given coverage-level threshold (horizontal axis) in each chromosome. Each base-position’s coverage-level was normalized by dividing it by the number of millions of read-bases mapped to that chromosome, separately for exonic and non-exonic bases. Six independent replicates of control (ethanol-treated, black lines) and MOG-7 AID (auxin-treated for 1h, blue lines) treatment groups were plotted. Dashed black boxes highlight exonic and non-exonic counts at the lower coverage threshold region of the plot, where the difference between treated and untreated samples in terms of non-exonic base-position coverage was most apparent. (D) Violin plot showing relative intronic read counts (vertical axis) vs expression level of host genes (horizontal axis). Black: Control (ethanol-treated); blue: MOG-7 AID (1h auxin treatment). (E) Visualization of splicing analysis of RNA-sequencing data comparing control (ethanol-treated) and MOG-7 AID (1h auxin treatment). Tracks are shown for selected genes *cls-2* and *klp-7* and are representative of all 6 replicates per condition. Red arrows indicate retained introns and red lines indicate the amplicon for RT-PCR validation of intron retention shown in (F). (F) RT-PCR validation of retained introns for *cls-2* (exon 4-5 PCR) and *klp-7* (exon 1-2 PCR) after MOG-7 AID for 1h and 2h. Position of the PCR primers used are shown in (E) in red beneath schematic showing gene structure.

We first examined exonic sequences to identify differentially expressed genes (DEGs) (Fig. S5B-D and Table S2). There were 243 and 702 DEGs (FDR < 0.05, absolute logFC > 0.585) identified after 1h and 2h of MOG-7 AID, respectively, with an overlap of 182 genes. Many of the DEGs identified were not germline-expressed genes (45% of 1h MOG-7 AID genes and 34% of 2h MOG-7 AID genes) (Fig. S5B-D). As we performed germline-specific MOG-7 AID, we supposed somatic DEGs may be responding to the auxin treatment rather than MOG-7 depletion. We subjected these DEGs to gene ontology (GO) overrepresentation analysis according to classification by *molecular function* and *biological process* using DAVID (https://david.ncifcrf.gov/). We detected an overrepresentation of GO terms related to xenobiotic stimuli and detoxification pathways, likely indicating responses to auxin exposure (Fig. S5E-H and Table S2). However, we also found overrepresentation of GO terms relating to meiotic chromosome separation, DNA damage and transcription in the 2h MOG-7 AID dataset, suggesting that these genes are responding to MOG-7 depletion and/or the associated germline defects (Fig. S5G-H).

Due to the predicted function for the MOG-7 protein domains in pre-mRNA splicing, we assessed the effect of germline-specific MOG-7 AID on intron retention (Fig. 4, Fig. S5 and Table S3). Focusing on the 1h MOG-7 AID dataset, we identified 5014 genes with retained introns, 93% of which are germline-expressed (Fig. 4B). We found that MOG-7 AID increased the global proportion of non-exonic RNA (intronic/intergenic) from autosomes (Fig. 4C and Fig. S6). The fact that the greatest difference was observed in low-coverage loci (less than 0.001 per million mapped bases) indicates that most of these differentially expressed regions are not unannotated exons but rather bona-fide introns or intergenic loci. Global changes in the proportion of non-exonic and exonic RNA were not observed for genes located on the X chromosome, probably due to the low level of transcription on the X chromosome in the germ line ^30^. We next examined intron reads per gene compared to expression level and found that as one may predict, genes with more transcripts have more remaining introns (Fig. 4D and Fig. S5B). We randomly selected two germline-expressed genes (*cls-2* and *klp-7*) to visualize intron retention caused by MOG-7 AID at specific loci (Fig. 4E). We found that all introns of these genes exhibit a degree of intron retention following 1h of MOG-7 AID (Fig. 4E). We validated the retention of a specific intron at each of these loci by PCR from independently isolated RNA samples for 1h and 2h MOG-7 AID (Fig. 4F). Together, these data show that acute loss of MOG-7 from the germ line causes widespread deficits in pre-mRNA splicing, that likely cause germ cell behavioral defects and germline sterility.

### The germ line recovers functionality when MOG-7 expression is restored

We have shown that germline depletion of MOG-7 causes developmental delay, sterility, failure of the sperm-oocyte switch and attenuation of cellular processes such as DNA synthesis, cell cycle progression, proliferation and nuclear organization (Figs. 1–2). These striking phenotypes are consistent with the splicing defects observed in thousands of germline-expressed genes when MOG-7 is depleted (Fig. 4).

Our germ cell analysis identified a cohort of arrested germ cells following long-term MOG-7 depletion (Fig. 2). We therefore wondered whether the germ line has the capacity to recover from such a severe breakdown in splicing and fundamental cellular processes. To examine this question, we used the flexibility of the AID system to monitor germline development after resupplying MOG-7 expression in adults. A previous study showed that protein expression recovers after auxin removal, with the rate of recovery dependent on the concentration of auxin used and potentially the developmental stage of auxin treatment ^23^. Further, protein recovery dynamics are dependent on gene-specific transcription and translation rates. To examine the dynamics of MOG-7::degron::GFP expression, we imaged hermaphrodite germ lines after 1h of auxin exposure and again 1-2h after auxin removal (Fig. S7). We found that MOG-7::degron::GFP expression was undetectable in the germ line after 1h of auxin exposure and robust expression was restored within the first hour of removal from auxin plates (Fig. S7).

Based on the ability of MOG-7 expression to rapidly recover following auxin removal, we established a protocol to examine the effect of restoring MOG-7 expression on germline function (Fig. 5A). We performed germline-specific MOG-7 AID for 18h from the early L4 stage until the first day of adulthood and then recovered animals on auxin-negative plates (Fig. 5A-B). We compared the brood size of these animals to those exposed to continual germline-specific MOG-7 AID and control animals (ethanol-treated for the same period) (Fig. 5B). We found that as shown in Figure 1G, animals exposed to continual MOG-7 AID were sterile (Fig. 5B). When we examined the broods of animals in recovery following MOG-7 AID for 18h, we observed few offspring during the first two days of adulthood (Fig. 5B). However, progeny production significantly recovered in 3-day adults and recovered hermaphrodites produced more progeny than control animals between days 4-6 (Fig. 5B). In total, MOG-7 AID recovered hermaphrodites generated a mean of 201 viable progeny (SD = 54.79) and 22 dead embryos (SD = 12), compared to control animals that generated a mean of 324 viable progeny (SD = 16.45) and 3 dead embryos (SD = 2). We wondered whether surviving progeny of MOG-7 AID recovered hermaphrodites were viable and fertile. From 56 randomly selected surviving L1 progeny of MOG-7 AID recovered adults, 54 developed into fertile adults, 1 was sterile and 1 was male. These data show that the germ line can functionally recover following restoration of MOG-7 expression.

**Figure 5.**
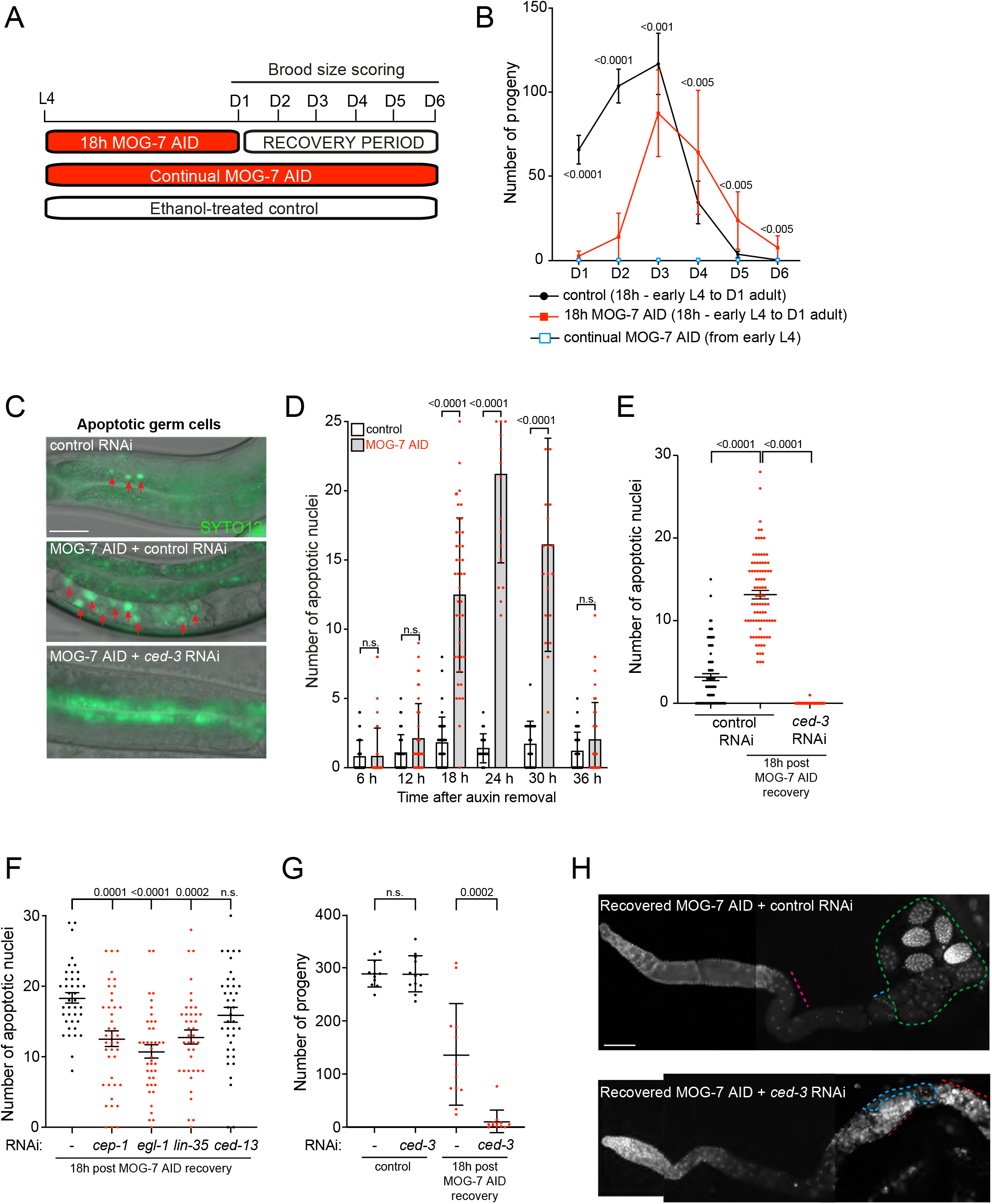
Apoptosis is required for recovery from MOG-7 depletion. In all experiments in this Figure, *mog-7::degron::gfp; sun-1p::TIR1::mRuby* hermaphrodites were used. (A) Experimental outline of MOG-7 depletion and recovery experiments. Early L4s were incubated on auxin plates for 18h after which animals were transferred to plates without auxin (MOG-7::degron:::GFP was detected within 1hr of auxin removal - Fig. S7) and brood size scored every 24h. Control animals were either continually exposed to auxin (continual MOG-7 AID) or control (ethanol-treated) from the L4 stage. (B) Quantification of daily brood size of control (black line), continual MOG-7 AID (blue box) or comparing 18h MOG-7 AID (red line). Broods were scored on days 1 to 6 following removal of auxin. Data expressed as mean ± S.D. and statistical significance was assessed by multiple t-tests with Holm-Sidak correction. n = 9-10. (C-D) Fluorescence micrographs (C) and quantification (D) of germ cell apoptosis (SYTO-12 staining). SYTO-12-positive nuclei were counted at hours 6, 12, 18, 24, 30 and 36 following 18h MOG-7 AID (auxin removal). Data expressed as mean ± s.e.m. and statistical significance was assessed by one-way ANOVA. n = 40 per time-point. Scale bar = 20 μm. (E) Quantification of germ cell apoptosis (SYTO-12 staining) +/- *ced-3* RNAi. SYTO-12 positive nuclei were counted 18h following removal of auxin. Data expressed as mean ± s.e.m. and statistical significance was assessed by one-way ANOVA multiple comparison with Sidak correction. n = 75. (F) Quantification of germ cell apoptosis (SYTO-12 staining) +/- *cep-1, egl-1, lin-35* or *ced-13* RNAi. SYTO-12 positive nuclei were counted 18h following removal of auxin. Data expressed as mean ± s.e.m. and statistical significance was assessed by one-way ANOVA multiple comparison with Sidak correction. n = 40. (G) Quantification of total brood of exposed hermaphrodites +/- *ced-3* RNAi. Data expressed as mean ± s.e.m. and statistical significance was assessed by unpaired t-test. n = 9-10. (H) Fluorescence micrographs of exposed hermaphrodites +/- *ced-3* RNAi. Germlines were extruded 48h after auxin removal. Germ lines stained with DAPI to visualize DNA (white). Scale bar = 25 μm. Upper panel - pink line = oocytes, blue line spermatheca, green dashed area = developing embryos. Lower panel - blue dashed area = misplaced sperm, red line = endomitotic nuclei/defective germ cells.

How is the adult germ line able to functionally recover from the extensive molecular and cellular deficits caused by MOG-7 depletion? We have shown that few offspring are generated in the first two days after restoring MOG-7 expression (Fig. 5B). As it takes approximately two days for PZ germ cells to develop into oocytes ^31^, we reasoned that this period may be utilized to remove defective germ cells from the germline. Indeed, programmed cell death (apoptosis) was previously shown to be required for the regenerative capacity of the germ line following extended starvation ^32^. To explore this hypothesis, we performed MOG-7 AID for 18h from early L4 larvae to young adult, and quantified apoptosis by SYTO-12 staining for 36h at 6-h intervals following auxin removal (Fig. 5C-D). We observed a large increase in the number of apoptotic cells between 18 and 30h after auxin removal, with a peak increase of 20-fold compared to control animals at the 24h time-point (Fig. 5D). Apoptosis functions as part of the physiological oogenesis program to control germ cell number and also facilitates germ cell death in response to DNA damage, environmental stress and pathogen exposure through distinct molecular pathways ^4–7^. These pathways converge at the CED-3 executioner caspase that is essential for germline apoptosis ^4^. We found that *ced-3* RNAi abrogates the burst of apoptosis in animals recovering from MOG-7 depletion (Fig. 5E). Conserved proteins acting upstream of CED-3 regulate physiological (LIN-35/Retinoblastoma) and DNA damage-induced (CEP-1/p53) apoptosis ^33, 34^. Two BH3-only EGL-1 and CED-13 proteins also act downstream of CEP-1 to promote DNA damage-induced apoptosis through partially parallel mechanisms ^35^. We used RNAi to knockdown expression of these genes and found that *cep-1, egl-1* and *lin-35* but not *ced-13* are partially required for germ cell apoptosis in MOG-7 AID recovered germlines (Fig. 5F). This suggests that apoptotic removal of defective germ cells in MOG-7 depleted germ lines is triggered by partially parallel pathways.

The data above suggest that apoptotic clearance of defective germ cells is a prerequisite for functional recovery of the germ line. We therefore measured the brood size of *ced-3* RNAi knockdown hermaphrodites following recovery from MOG-7 depletion (Fig. 5G). *ced-3* knockdown abolished the ability of the germ line to recover from MOG-7 AID, whereas *ced-3* RNAi animals exposed to ethanol had indistinguishable broods from RNAi control animals (Fig. 5G). DAPI staining of extruded germ lines 48h after MOG-7 AID recovery detected oocytes and multiple developing embryos in control RNAi animals (Fig. 5H - top panel). In contrast, germ lines of *ced-3* RNAi animals were severely disorganized with many misplaced sperm and masses of proximal DAPI staining, likely representing endomitotic nuclei (Fig. 5H - bottom panel). Together, these data reveal that the removal of defective cells by apoptosis is essential for the *C. elegans* germ line to recover following a breakdown in RNA splicing.

### Functional gametes are generated from distal progenitors after MOG-7 recovery

What is the source of viable gametes that enables germline recovery upon restoring MOG-7 expression? We reasoned that these gametes either originate from arrested/slow cycling germ cells in the progenitor zone that re-enter the cell cycle or that meiotic germ cells can resolve the molecular and cellular deficits caused by MOG-7 depletion. To distinguish these scenarios, we used EdU to label germ cells generated in the progenitor zone during MOG-7 depletion (in-auxin EdU) or after MOG-7 expression was restored (post-auxin EdU) (Fig. 6A). In-auxin EdU was applied 2h after commencing auxin treatment to ensure that EdU-labelled germ cells are MOG-7 depleted (Fig. 6A).

**Figure 6.**
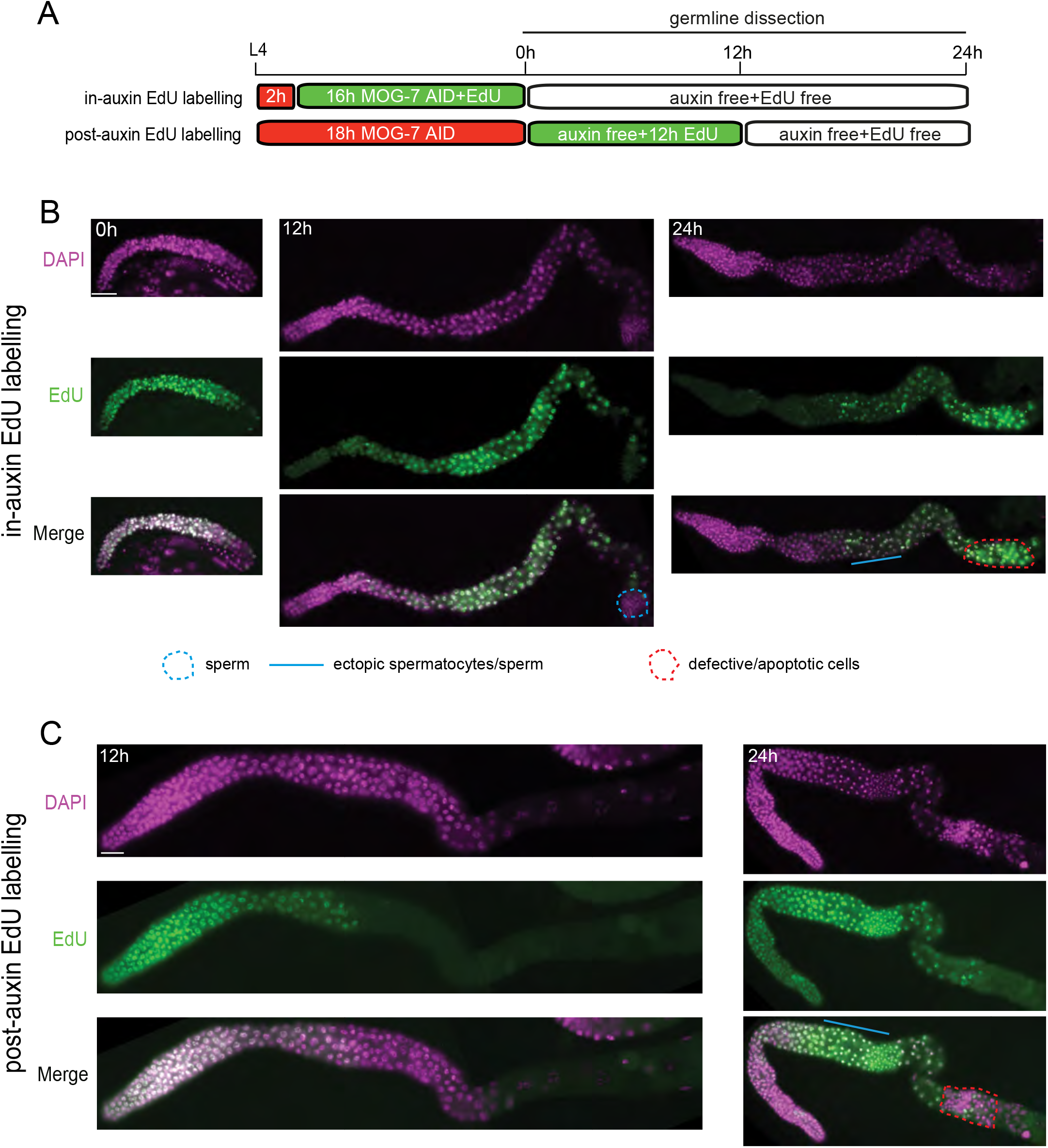
New gametes are generated after MOG-7 is resupplied. (A) Experimental outline for S-phase labelling of adult germ cells following *mog-7* depletion and recovery. Early L4 *mog-7::degron::gfp; sun-1p::TIR1::mRuby* hermaphrodites were incubated as follows: in-auxin labelling (2h auxin, 16h auxin + EdU, 24h auxin free and EdU free); post-auxin EdU labelling (18h auxin, 12h auxin free and 12h EdU, 12h auxin free and EdU free). Germ lines were extruded and analyzed at 0h, 12h and 24h following the 18h of MOG-7 depletion in both labelling methods. (B-C) Fluorescence micrographs of germ lines exposed to in-auxin EdU labelling (B) and post-auxin EdU labelling (C). Germ lines stained with DAPI to visualize DNA (magenta) and labelled with EdU to detect replicated DNA (green). Ectopic spermatocytes/sperm = blue line/dashed line. Defective/apoptotic cells = red dashed area. Scale bar = 20μm.

After 16h of in-auxin EdU labelling, most germ cells are EdU^+^, confirming cell cycle progression occurs during MOG-7 depletion (Fig. 6B - 0h time-point). Previously, we identified a cohort of arrested germ cells in MOG-7 depleted germ lines (Fig. 2H and J). We hypothesized that these germ cells may re-enter the cell cycle to replenish the germ line after restoring MOG-7 expression. Consistent with this, after removing auxin and EdU, we observed many distal EdU^-^ germ cells, suggesting that these cells re-entered the cell cycle after restoring MOG-7 expression (Fig. 6B - 12-24h time-points). Germ cells that are labelled with EdU 24h after auxin/EdU removal are mostly ectopic sperm/spermatocytes, oocytes, and germ cells with highly condensed proximal nuclei that are likely cleared by apoptosis (Fig. 6B - 24h time-point). These EdU^+^ cells could have been generated during MOG-7 depletion or while MOG-7 expression was being restored, but likely have irreparable splicing defects due to the extended loss of MOG-7.

In post-auxin EdU labelling, ectopic sperm/spermatocytes (Fig. 6C - blue line in 24h time-point) and a small number of highly condensed proximal nuclei (Fig. 6C - red dashed box in 24h time-point) are EdU^+^, indicating that they were labelled after auxin removal. However, most of these highly condensed proximal nuclei are EdU^-^, suggesting they were generated during MOG-7 depletion and exited the cell cycle prior to auxin removal (Fig. 6C - red dashed box in 24h time-point). These data reveal that 1) germ cells entering meiosis during MOG-7 depletion are unable to functionally recover and are likely cleared by apoptosis, and 2) that the initial cohort of germ cells generated after restoring MOG-7 expression acquire the sperm fate. In support of this, ~60% of germlines are spermatogenic 24-36h after auxin removal, with the oogenic program resuming after 48h (Fig. S8). Together, these data suggest that germline recovery requires the generation of new germ cells in the progenitor zone. To independently verify this hypothesis, we used RNAi to knockdown expression of CDK-1, a cyclin-dependent kinase required for mitotic divisions ^36^. We found that germline recovery from MOG-7 depletion requires CDK-1 function (Fig. S9).

## DISCUSSION

In this study, we identified a critical function for the previously uncharacterized protein MOG-7 in the *C. elegans* hermaphrodite germ line. We found that germline-specific loss of MOG-7 causes fully penetrant sterility. We observed multiple germ cell morphology and cell cycle defects in MOG-7 depleted germ lines that likely prevent the generation of viable gametes. Detection of *in vivo* MOG-7-interacting proteins identified multiple pre-mRNA splicing factors with known roles in spliceosome disassembly. To support the function of MOG-7 in pre-mRNA splicing, we found that MOG-7 is important for removal of introns from over 5000 proteincoding genes in the germ line. Despite being encumbered by extensive cellular and molecular deficits, hermaphrodites can become fertile within hours of restoring MOG-7 expression. As part of its recovery, the germ line undergoes a burst of apoptotic death, presumably to remove defective germ cells and to provide recycled cytoplasmic components for a newly generated germ cell cohort. Indeed, if apoptosis is inhibited MOG-7-recovered animals are sterile. We detected a slowing of the cell cycle and germ cell arrest in the distal germ line following long-term MOG-7 depletion. Collectively, these cells likely provide the source of fertile gametes once MOG-7 expression and correct pre-mRNA splicing is restored to the germ line. Together, these observations highlight the remarkable plasticity of the germ line and its ability to recover from extensive defects in fundamental cellular and molecular functions.

### MOG-7 is an orthologous to yeast and human splicing factors

In *S. cerevisiae*, the Nineteen complex-related proteins Ntr1 and Ntr2 are critical regulators of spliceosome disassembly ^15, 17^. Ntr2 promotes the recruitment of Ntr1 to the intron lariat spliceosome complex. Ntr1 then binds and activates Prp43 which induces spliceosome disassembly, where spliced introns are degraded and spliceosomal proteins recycled. The human genome does not encode a recognizable Ntr2 ortholog. However, the C2orf3 protein, which contains a GC-rich sequence DNA-binding factor (GCFC) domain, interacts with the human ortholog of Ntr1 (TFIP11), in addition to Prp43, and is involved in intron turnover ^18^. The *mog-7* locus encodes two proteins related to Ntr2 (*S. cerevisiae*) and GCFC (human). MOG-7a contains both an Ntr2 and GCFC domain, whereas MOG-7b contains only the GCFC domain. Our experiments show that removal of both MOG-7 isoforms causes intron retention, germ cell defects and sterility. Further, both MOG-7a and MOG-7b interact with the *C. elegans* spliceosome components we identified by mass spectrometry. Our *in vitro* immunoprecipitation experiments showed that both the Ntr2 and GCFC domains of MOG-7 are able to interact with spliceosome proteins. In other organisms, the Ntr2 and GCFC domains are present in distinct proteins ^15, 18^. Whether the MOG-7a (Ntr2 and GCFC) and MOG-7b (GCFC only) isoforms perform cell-specific or spliceosome complex-specific roles is unknown, and is an interesting avenue of future investigation.

### Loss of MOG-7 rapidly causes intron retention

Spliceosomes form on each newly synthesized intron before disassembling for the next round of splicing ^37^. In a highly ordered process, spliceosome sub-complexes sequentially associate with each intron in a reversible manner ^38^. Hence, an interruption in spliceosome protein disassembly inhibits the splicing of subsequent transcripts. We found that MOG-7 directly interacts with multiple splicing factors involved with spliceosome disassembly and observed extensive intron retention within 1h of depleting the MOG-7 protein. Such rapid retention of introns suggests that the splicing process is highly sensitive to MOG-7 removal and that perhaps introns are retained even as MOG-7 protein levels start to reduce following auxin treatment. We have shown that restoring MOG-7 expression following long-term MOG-7 depletion enables animals to become fertile. Whether restoring MOG-7 expression enables spliceosomes to reform and initiate splicing on incorrectly spliced transcripts or whether new transcription is required for recovery is unknown. The requirement for apoptosis in the recovery process (see below) indicates that many germ cells, and their defective transcripts, need to be removed to enable the generation of functional gametes. However, there may be a cohort of germ cells that can functionally recover from the splicing defects.

### MOG-7 depletion affects germ cell morphology and cell cycle progression

A previous study showed that starving adult hermaphrodites causes cell-cycle arrest where germ cells stop dividing and become quiescent ^39^. These germ cells exhibit a slowing of S-phase and arrest in G2 of the cell cycle. Upon re-feeding these quiescent cells rapidly re-enter M-phase and produce viable progeny ^39^. Our analysis of the germ line following MOG-7 depletion revealed multiple defects in germ cell nuclear morphology, clustering of germ cells and areas of the germ line devoid of germ cells. In addition, the progenitor zones of MOG-7 depleted animals have a reduced number of germ cells in M-phase and S-phase of the cell cycle. These findings suggest a slowing of the germ cell cycle in the progenitor zone in response to the splicing defects. Examination of germ lines following long-term MOG-7 depletion also revealed a number of arrested germ cells (Fig. 2) that are likely a source of new gametes.

### Apoptotic disposal of germ cells enables recovery from MOG-7 depletion

In the hermaphrodite germ line, both physiological and stress-induced germ cell death require induction of the core apoptotic caspase CED-3 ^4, 6^. When we re-supplied MOG-7 following MOG-7 depletion, the germ line exhibited a burst of apoptosis that was dependent on CED-3. Initiation of apoptosis did not occur until 18h after MOG-7 expression was restored. This delay may be caused by the defective splicing of genes required for apoptosis in MOG-7-depleted germ lines, such that these mis-spliced transcripts need to be resolved and/or new transcription activated prior to initiation of apoptosis. In support of this premise, transcripts of the core apoptotic pathway genes (*ced-3*, *ced-4*, *ced-9*) contain retained introns within 1h of MOG-7 depletion (Table S3). We showed that *ced-3* knockdown caused sterility in MOG-7-recovered but in not control animals. Indeed, the proximal end of MOG-7-recovered *ced-3* knockdown germ lines is replete with aberrant germ cells. This confirms the importance of removing defective germ cells by apoptosis for germline recovery after resupplying MOG-7 expression. Parallel pathways act upstream of CED-3 to control physiological (LIN-35/Retinoblastoma) and stress-induced germ cell death (CEP-1/p53, EGL-1/BH3-only and CED-13/BH3-only ^33–35^. We found that LIN-35, CEP-1 and EGL-1 but not CED-13 are partially required for apoptotic induction in MOG-7 AID recovered germlines. The role of each cell death pathway in this context may be dependent on the nature of the germ cell defect and/or the stage of the germ cell cycle at which splicing collapse occurs.

### Regulation of sperm/oocyte cell fate

Multiple studies have previously revealed that downregulation of splicing factors cause inappropriate masculinization of the germ line (Mog phenotype) ^8, 9, 12, 13, 21, 40, 41^. Indeed, *F43G9.12/mog-7* RNAi knockdown was previously shown to cause synthetic germline masculinization and lethality in mutants for the PUF domain RNA-binding proteins FBF-1 and FBF-2 ^9^. These FBF proteins repress genes that promote differentiation, meiotic processes and genes associated with spermatogenesis ^42–44^. This study also found that RNAi knockdown of *F43G9.12/mog-7* alone resulted in a low penetrant Mog phenotype ^9^. We took advantage of the auxin-inducible degron system to completely remove F43G9.12/MOG-7 protein at different developmental time-points. We revealed that removing MOG-7 at the L3 larval stage caused masculinization of most germ lines, whereas removing MOG-7 from the L4 stage resulted in mostly oogenic germ lines. This suggests the importance of splicing regulation prior to/during the sperm-oocyte switch.

When examining germline behavior during recovery from MOG-7 depletion, we identified an interesting switch in germ line sexual fates. As mentioned previously, continuous MOG-7 AID from L4 resulted in mostly oogenic germline development. Indeed, when we depleted MOG-7 from early L4 to the first day of adulthood, we initially observed oogenic germ lines. However, 24 and 36h after restoring MOG-7 expression the germ lines were spermatogenic, and then reverted to oogenesis 12h later. The initial oogenic phase during MOG-7 recovery likely enables the apoptotic program (oogenic-specific) to remove defective germ cells ^4^. After the defective germ cells are removed, the germ line then re-enters a new phase of spermatogenesis prior to generating oocytes. These findings highlight the plasticity of the germ line and perhaps the sensitivity of cells at different stages of the cell cycle and differentiation process to a disruption in splicing.

Together, this study reveals the ability of the *C. elegans* germ line to recover functionality following a breakdown in pre-mRNA splicing. Removal of the MOG-7 splicing factor induces multiple defects in germ cell morphology and behavior throughout the germ line that leads to sterility. After restoring MOG-7 expression in adults, the germ line initiates apoptotic clearance of defective germ cells and reinitiation of germ cell proliferation.

## SUPPLEMENTARY MATERIAL

### SUPPLEMENTARY FIGURES

**Figure S1. *F43G9.12/mog-7* RNAi experiments**

(A) Quantification of one-day adult mutant germ lines +/- *mog-7* RNAi from the L1 larval stage. Data expressed as mean ± s.e.m. of three replicates (n = 25 per replicate) and statistical significance was assessed by unpaired t-test.

(B-C) Quantification of developmental success (LL - live larvae vs DE - dead eggs) of hermaphrodites +/- *mog-7* RNAi from the L1 (B) or L4 (C) larval stage. RNAi experiments were performed in *rrf-1(pk1417)* mutant animals. Data expressed as mean ± s.e.m. and statistical significance was assessed by unpaired t-test.

**Figure S2. *mog-7::degron::GFP* expression analysis**

(A-E) DIC and fluorescence micrographs of *mog-7::degron::gfp* expression. (A) Embryonic expression. Scale bar = 20 μm. (B) Head expression (L1 larva). Scale bar = 20 μm. (C) Tail expression (L1 larva). Scale bar = 20 μm. (D) Vulval expression (early L4 - left; late L4 - right). Scale bar = 5μm. (E) L1 gonad - Z1/Z4 = somatic gonad precursors and Z2/Z3 = primordial germ cells. Scale bar = 5 μm.

(F) Western blot of *mog-7::degron::gfp* expression detected with anti-GFP antibody. Predicted molecular weights = 121kD (MOG-7a::degron::GFP) and 62kD (MOG-7b::degron::GFP).

**Figure S3. Control experiments for the *mog-7::degron::GFP* strain**

(A) Quantification of germ cell nuclei number in wild-type and *mog-7::degron::GFP* adult hermaphrodites. Data expressed as mean ± s.e.m. and statistical significance was assessed by multiple t-test (Holm-Sidak correction). n = 5-6. n.s. not significant.

(B) Brood size of wild-type, *mog-7::degron::GFP* and *mog-7::degron::GFP; sun-1p::TIR1::mRuby* hermaphrodites. Data expressed as mean ± s.e.m. and statistical significance was assessed by ordinary one-way ANOVA. n = 3-5. n.s. not significant.

(C) Quantification of germ cell nuclei number in wild-type and *mog-7::degron::GFP* adult hermaphrodites (+/- auxin). Data expressed as mean ± s.e.m. and statistical significance was assessed by multiple t-test (Holm-Sidak correction). n = 5-6. n.s. not significant.

(D) Quantification of germ cell nuclei number in wild-type and *sun-1p::TIR1::mRuby* adult hermaphrodites (+/- auxin). Data expressed as mean ± s.e.m. and statistical significance was assessed by multiple t-test (Holm-Sidak correction). n = 6-9. n.s. not significant.

(E) Fluorescence micrographs of *mog-7::degron::gfp* expression in the head of *mog-7::degron::GFP; sun-1p::TIR1::mRuby* adult hermaphrodites exposed to ethanol (left) or 1mM auxin (right) for 1h. Scale bar = 10 μm.

**Figure S4. Input Western blots for co-immunoprecipitation experiments**

(A) Immunoprecipitation of MOG-7a and MOG-7b followed by Western blot in COS-7 cells confirming the interaction of both MOG-7 isoforms with spliceosome proteins (tagged with a FLAG epitope) identified by mass spectrometry (n = >2 independent experiments). Blot shows MOG-7a and MOG-7b in each sample detected with anti-V5 antibody. Inputs are shown in (B), where each interacting protein is detected with an anti-FLAG antibody.

(C) Immunoprecipitation of MOG-7(ΔGCFC) followed by Western blot in COS-7 cells confirming its interaction with spliceosome proteins identified by mass spectrometry. Inputs are shown in (D).

**Figure S5. RNA sequencing analysis**

(A) Multi-dimensional scaling (MDS) plot of moderated gene-wise counts per million (CPM) values, only counting canonical exon-mapped reads. Shown are ethanol-treated controls (black) and 1 h (blue) or 2h (red) auxin-treated MOG-7 AID animals with six independent replicates per condition. Analysis was performed using Limma-voom (Law et al., 2014) and visualized with Degust.

(B) Summary of the number of differentially-expressed genes (DEGs; genes having an FDR of < 0.05 and absolute logFC > 0.585) after germline-specific MOG-7 AID for 1h and 2h compared to ethanol-treated control. Number of DEGs in each condition shown in parentheses. Venn diagram depicts the percentage of DEGs common to both time-points (75%) and those DEGs only identified after 2h.

(C-D) Summary of the number of differentially-expressed genes (DEGs) after germline-specific MOG-7 AID for 1h (C) and 2h (D) compared to ethanol-treated control. Number of DEGs in the germ line (GL) or non-germ line (non-GL) shown in parentheses.

(E-H) Gene ontology (GO) term enrichment for biological process using DAVID (https://david.ncifcrf.gov/), on DEGs identified by contrasting the control vs. 1h auxin-treated (E-F), and control vs. 2h auxin-treated (G-H), MOG-7 AID gene-wise exonic read-counts from the RNA-seq experiments. The *biological process* (E, G) and *molecular function* (F, H) gene ontology classes were searched for terms enriched in up- and down-regulated DEGs.

**Figure S6. RNA splicing analysis**

(A) Percentage of exonic and non-exonic base-positions (vertical axis) exceeding any given coverage-level threshold (horizontal axis) in each chromosome. Each base-position’s coverage-level was normalized by dividing it by the number of millions of read-bases mapped to that chromosome, separately for exonic and non-exonic bases. Six independent replicates of control (ethanol-treated, black lines) and MOG-7 AID (auxin-treated for 2h, red lines) treatment groups were plotted. Dashed black boxes highlight exonic and non-exonic counts at the lower coverage threshold region of the plot, where the difference between treated and untreated samples in terms of non-exonic base-position coverage was most apparent.

(B) Violin plot showing relative intronic read counts (vertical axis) vs expression level of host genes (horizontal axis). Black: Control (ethanol-treated); red: MOG-7 AID (2h auxin treatment).

**Figure S7. Qualitative analysis of *mog-7::degron::gfp* depletion/recovery**

DIC and fluorescence micrographs of *mog-7::degron::gfp* expression. 18h after incubation on auxin plates (MOG-7 expression depleted in the germ line - upper two panels); 1h and 2h after transfer from auxin to NGM plates (MOG-7 expression restored in the germ line - center and lower panels). Dashed line = germ line region. Scale bar = 10 μm.

**Figure S8. Analysis of germ line sex following MOG-7 depletion and recovery**

(A) Experimental outline of *mog-7* depletion and recovery experiments. Early L4 *mog-7::degron::gfp; sun-1p::TIR1::mRuby* hermaphrodites were incubated on 0.1mM auxin plates for 18h after which animals were transferred to plates without auxin and spermatogenic/oogenic fate scored 12h, 24h, 36h, 48h and 72h following auxin removal. Control animals were either continually exposed to auxin (Continual MOG-7 AID) or ethanol (control) from the L4 stage.

(B-C) Quantification (B) and fluorescence micrographs of germ lines (C) of *mog-7::degron::gfp; sun-1p::TIR1::mRuby* hermaphrodites exposed to ethanol, MOG-7 AID from the early L4 stage for 18h or continual MOG-7 AID. The percentage of masculinized germ lines are shown in B. Data expressed as mean ± s.e.m. and statistical significance was assessed by multiple t-tests with Holm-Sidak correction. n = 27-50 for each replicate. Germ lines stained with DAPI to visualize DNA (white). Scale bar = 10 μm.

**Figure S9. CDK-1 is required for germline recovery following MOG-7 depletion**

Quantification of total brood of control (L4440 vector) and *cdk-1* RNAi hermaphrodites following 18h of MOG-7 AID from the L4 stage (as in Fig 5A-B). Note: *cdk-1* RNAi progeny died during embryogenesis. Data expressed as mean ± s.e.m. and statistical significance was assessed by unpaired t-test. n = 11-12.

### TABLES

**Table S1. MOG-7 co-immunoprecipitation results**

Immunoprecipitation of MOG-7a and MOG-7b followed by Western blot in COS-7 cells confirming the interaction of both MOG-7 isoforms with spliceosome proteins identified by mass spectrometry (n = >2 independent experiments). Green box = interaction detected, grey box = no interaction detected, white box = not determined.

**Table S2. Differentially-expressed gene lists after MOG-7AID**

**Table S3. Genes contain increasing non-exonic reads after MOG-7AID**

**Table S4. Strains used in this study**

**Table S5. Plasmids used in this study**

**Table S6. Oligos used in this study**

## EXPERIMENTAL PROCEDURES

### Contact for reagent and resource sharing

Strains used in this study will be deposited at the *Caenorhabditis* Genetics Center (CGC) and will be available upon request. Further information and requests for resources and reagents should be directed to and will be fulfilled by the Lead Contact Roger Pocock.

### Experimental models and subject details

#### Caenorhabditis elegans

*C. elegans* strains were cultured on Nematode Growth Medium (NGM) plates and fed with OP50 *Escherichia coli* bacteria at 20°C, unless otherwise stated. All strains used in this study are listed in the Table S4. Experiments were performed in triplicates and the number of animals analyzed is annotated in each figure and legend where appropriate.

#### Mammalian cells

For co-immunoprecipitation experiments, COS-7 cells were cultured in Dulbecco’s Modified Eagle Medium (containing 4500mg/L d-glucose, 584mg/L L-glutamine, 110mg/L sodium pyruvate) (Thermo Fisher) containing 1% penicillin and streptomycin (Invitrogen), 5μM L-glutamine (Life Technologies) and 5% fetal bovine serum (Thermo Fisher).

### Endogenous tagging of MOG-7 with degron::GFP using CRISPR-Cas9

A C-terminal MOG-7::degron::GFP knock-in strain was generated using CRISPR/Cas9-triggered homologous recombination ^22^. The crRNA (AGTTTTTGGAGAAACTCGAG) was designed using the online tool provided by https://sg.idtdna.com. Asymmetric-hybrid donor repair templates were generated by PCR using ultramer primers (see primer list). The following mix was then injected into wild-type animals: 4μg repair template, 5μg Cas9 protein, 2μg universal tracrRNA, 1.1μg crRNA, *myo-2::mCherry* plasmid (4ng/μl). Individual F1 progeny of injected wild-type worms were picked to individual plates and F2 progeny screened for *degron::gfp* knock-in by PCR. After confirmation of insertion by Sanger sequencing, the *degron::gfp* knock-in was outcrossed three times prior to analysis. MOG-7::degron::GFP was imaged using a Zeiss Axiocam 40x objective.

### RNA interference experiments

HT115(DE3) *E. coli* bacteria expressing RNAi plasmids for specific genes or empty vector (L4440) were grown in Luria Broth (LB) + Ampicillin (50μg/ml) at 37°C for 16hr. Saturated cultures of RNAi bacteria were plated on RNAi plates and allowed to dry for 24hr. Hermaphrodite larvae were placed on the RNAi plates and incubated for various periods depending on the experiment (described in each figure legend).

### Auxin-inducible degradation (AID) plates

NGM plates were supplemented with 0.1mM or 1mM Auxin (indole-3-acetic acid, Alfa Aesar, ALFA10556) from a stock of 400mM dissolved in ethanol. NGM plates supplemented with ethanol were used as a control.

### Brood size analysis

10 L4 hermaphrodites were picked onto individual NGM plates seeded with OP50 bacteria. Worms were allowed to lay eggs for 24hr and then the mothers were individually moved to new plates. After a further 24hr embryos were analyzed for hatching. This process was repeated for six days. The number of larvae and embryos were counted each day and summed as the total brood size.

### Transfection of COS-7 cells

60,000 cells were transferred into each well of a 24-well plate. Plasmid transfection was performed when cells were approximately 70% confluent. In each well, 0.25μl of plasmid was mixed with 70μl Opti-MEM®, and 2.5μl lipofectamine was mixed with 70μl Opti-MEM®. After incubating at room temperature for 5min, the two mixtures were combined and incubated at room temperature for 20 min. The 140μl transfection complexes were applied to each well. After incubating in 37°C for 3h, 1 ml of fresh media was added to each well and cells were harvested 1 day after the transfection.

### Cell lysis and co-immunoprecipitation

Each cell culture well was washed 3 times with cold 1x PBS buffer before 100μl cell lysis buffer (containing 50mM Tris pH 7.6, 150mM NaCl, 1% TX-100, cOmplete™ EDTA-free Protease Inhibitor Cocktail) was added. Cells were scraped off the bottom of the plate and sonicated (10 sec on, 20 sec off, 3 cycles) before being centrifuged at the highest speed for 10 min. 20μl of the supernatant was extracted as the input.

1μl Mouse anti-V5 antibody was added to every 10μl Dynabeads™ protein G (Thermo Fisher Scientific) (2.5μl per sample) and left to incubate overnight to facilitate binding before use. Crosslinking was performed by incubating the antibody-coupled Dynabeads™ in 250μl 5mM Pierce BS3 Crosslinker (in PBS buffer) rotating for 30 min at room temperature. The crosslinking reaction was then quenched by adding 12.5μl 1M Tris HCl (pH 8). After washing thrice with 1x PBS buffer, the crosslinked Ab-Dynabeads™ were added into the supernatant of cell lysates for immunoprecipitation. After a 2h incubation, the Dynabeads™ were isolated using a magnet, and washed thrice with cell lysis buffer.

### Western blotting

#### COS-7 samples

After mixing with 1xBolt™ LDS sample buffer (Invitrogen™) and 1xBolt™ Sample reducing Agent (Invitrogen™), the input sample and postimmunoprecipitation Dynabeads™ were boiled at 95°C for 5 min. 10% resolving gels were used to separate proteins, except for samples containing FLAG-PRP-8 (276 kD), in which case 6% resolving gels were used. The components of the gels are shown in Table 2.9. SDS-PAGE was performed in a Mini-PROTEAN Tetra Vertical Electrophoresis Cell (BIO-RAD) filled with 1x SDS-PAGE Running buffer (10x SDS-PAGE Running buffer: 30 g of Tris base, 144 g of glycine, and 10 g of SDS in 1000 ml of H2O). Voltage of 60V and 120V were respectively used for stacking and protein separation, respectively. After separation on an SDS-PAGE gel, proteins were transferred to the PVDF membrane by using iBlot™ 2 Transfer Stacks (Thermo Fisher Scientific) and iBlot™ 2 Gel Transfer Device (Thermo Fisher Scientific) according to the manufacturer’s instructions. After blotting, the PVDF membrane was blocked through incubation in 5% w/v bovine serum albumin (in 1x PBS) at room temperature for 30 min. Subsequently, the membrane was incubated in primary antibody at 4°C overnight, followed by an incubation with secondary antibodies at room temperature for 1.5h. Primary antibodies (Mouse anti-FLAG/DYKDDDDK tag antibody (2368, Cell Signaling), Mouse anti-V5 tag antibody (SV5-Pk1, Bio-Rad) and secondary antibodies (Goat anti-Mouse IgG (H+L) secondary antibody, HRP conjugate (32430, Thermo Fisher Scientific)) were diluted (1:1000) in 1% w/v bovine serum albumin (in 1x PBS). 1x PBS (containing 0.1% Tween-20) was used to wash the membrane after each antibody incubation. Proteins were detected using the EZ-ECL Chemiluminescence Detection Kit for HRP (Biological Industries) and ChemiDoc XRS+ Gel Documentation System (Bio-Rad Laboratories). Images were taken using the ImageLab™ software (Bio-Rad).

#### *C. elegans* samples

Transgenic worms expressing degron::GFP-tagged MOG-7 were grown on NGM plates coated with OP50 bacteria. A packed volume of 1ml of mixed stage worms and embryos were collected by washing with M9 buffer. Worms were washed three times with M9 buffer, pelleted by centrifugation and 3ml of lysis buffer (50mM Tris pH 7.4, 150mM NaCl, 2% triton X-100, 0.1% SDS, 1x protease inhibitor cocktail) was added. Worms were disrupted using a mortar and pestle, then sonicated using Bioruptor^®^ (Diagenode) at high amplitude (4°C; 30 sec ON and 20 sec OFF) for 10-15 cycles. After centrifuging at 3000rpm for 5min, the supernatant was collected and GFP-Trap® (Chromotek) was used to precipitate MOG-7::degron::GFP. Samples were cooled to room temperature and run on a 10% polyacrylamide gel. The gel was blotted to a PVDF membrane using the iBlot semidry blot system (Thermo Fisher). After blocking with 5% BSA in PBS for 1hr, the PVDF membranes were incubated with anti-GFP antibody (Roche) in 1% BSA in PBS-tween (0.1%) for 16hr at 4°C. The membrane was then washed three times with PBS-tween and incubated with horseradish peroxidase-conjugated secondary antibodies in 1% BSA in PBS-tween for 1hr at 22°C. The membrane was washed five times with PBS-tween and incubated with ECL reagents (Thermo Fisher) for 2min at room temperature. The membrane was exposed using Biorad ChemiDoc XRS+ and images taken using a CCD camera and Image lab 6.0.1 software.

### Mass spectrometry

Mixed-stage worms containing mostly one-day-old adults, L1s and embryos were harvested for mass spectrometry (MS) analysis. Approximately, 400,000 adults/per sample were required for robust MOG-7::degron::GFP pulldown. Three genotypes were analyzed in each MS experiment: *sun-1p::TIR1; mog-7::degron::gfp, evIs111* (GFP-only control), wild-type (no fluorophore control). Harvested worms were washed with M9 buffer (five times), before adding three times dry worm volume of RIPA buffer. After grinding and sonicating (30 sec on, 30 sec off, 30 cycles), each sample was incubated with a mixture of 10μl IgG conjugated magnetic beads (Cell Signaling) and 10μl Dynabeads™ for 1h to preclear. After removing the IgG conjugated magnetic beads, each sample was incubated with 10μl GFP-Trap® (Chromotek) at 4°C overnight for immunoprecipitation, and a *sun-1p::TIR1; mog-7::degron::gfp* sample was incubated with 10μl IgG conjugated magnetic beads (Cell Signaling) as a control.

Magnetic beads were collected using a magnet and washed with buffers in the following order: wash buffer 1 (RIPA buffer - 50mM Tris pH7.6, 150mM NaCl, 1% TX-100, 0.1% SDS), wash buffer 2 (50mM Tris pH 8.0, 150 mM NaCl), wash buffer 3 (50mM Tris pH 8.0, 450mM NaCl), and wash buffer 4 (50mM Tris pH 8.0). Then, beads were then incubated in 150μl 0.2 M glycine (pH 2.5) for 5 min to elute the protein. The elution was repeated thrice, and the combined 450μl of eluted samples were neutralized with ~80μl 1M Tris/HCl pH 8.0.

MS analysis was performed by Monash Biomedical Proteomics Facility. The Nano LC system used in this study was the Dionex Ultimate 3000 RSLCnano, the Mass spectrometer used was the QExactive Plus 2 (Thermo Scientific), the analytical column used was the Acclaim PepMap RSLC (75μm x 50cm, nanoViper, C18, 2μm, 100A□; Thermo Scientific), and the Trap column used was the Acclaim PepMap 100 (100μm x 2cm, nanoViper, C18, 5μm, 100A□; Thermo Scientific). The search engine Byonic (ProteinMetrics) was used for data analysis.

### Molecular cloning

For mammalian expression, the following plasmids were used to insert *C. elegans* cDNA sequences: pcDNA3.1(zeo) and p3XFLAG-CMV-10.

#### mog-7a cDNA::V5 tag

The *mog-7a cDNA::V5* mammalian expression construct was generated by cloning the 2430 bp *mog-7a* cDNA with HindIII into the pcDNA3.1(zeo) expression vector.

#### mog-7b cDNA::V5 tag

The *mog-7b cDNA::V5* mammalian expression construct was generated by cloning the 930 bp *mog-7b cDNA* with HindIII into the pcDNA3.1(zeo) expression vector.

#### *eftu-2cDNA::FLAG* tag

The *eftu-2cDNA::FLAG* mammalian expression construct was generated by cloning the xx bp *eftu-2* cDNA with NheI-HindIII into the p3XFLAG-CMV-10 expression vector.

#### *Y94H6A.3cDNA::FLAG* tag

The *Y94H6A.3cDNA::FLAG* mammalian expression construct was generated by cloning the 561 bp *Y94H6A.3* cDNA with KpnI into the p3XFLAG-CMV-10 expression vector.

#### *stip-1cDNA::FLAG* tag

The *stip-1cDNA::FLAG* mammalian expression construct was generated by cloning the 2493 bp *stip-1* cDNA with XbaI into the p3XFLAG-CMV-10 expression vector.

#### *prp-8cDNA::FLAG* tag

The *prp-8cDNA::FLAG* mammalian expression construct was generated by cloning the 6990 bp *prp-8* cDNA with XbaI into the p3XFLAG-CMV-10 expression vector.

#### *skp-1cDNA::FLAG* tag

The *skp-1cDNA::FLAG* mammalian expression construct was generated by cloning the 1608 bp skp-1 cDNA with XbaI into the p3XFLAG-CMV-10 expression vector.

#### *prp-19cDNA::FLAG* tag

The *prp-19cDNA::FLAG* mammalian expression construct was generated by cloning the 1930 bp *prp-19* cDNA with XbaI into the p3XFLAG-CMV-10 expression vector.

For RNAi experiments, the following plasmids were generated.

#### *cep-1* RNAi

The *cep-1* RNAi construct was generated by cloning the 571 bp *cep-1* genomic DNA (3824-4395) with HindIII into the L4440 vector.

#### *egl-1* RNAi

The *egl-1* RNAi construct was generated by cloning the 814 bp *egl-1* genomic DNA with HindIII into the L4440 vector.

#### *ced-13* RNAi

The *ced-13* RNAi construct was generated by cloning the 348 bp *ced-13* genomic DNA with HindIII into the L4440 vector.

### Germline analysis

Germline analysis was performed as previously reported ^46^. L4 hermaphrodites were picked to OP50 plates and incubated for 16hr at 20°C to reach the young adult stage. Germ lines were extruded from sedated worms and fixed on a poly-L-lysine coated slides using ice cold methanol for 1min and then in 3.7% paraformaldehyde (PFA) for 25min. Fixed germ lines were washed three times in phosphate buffered saline (PBS, pH 7.4) and blocked using 30% normal goat serum before incubating with primary antibodies overnight at 4°C. After incubation, germ lines were washed three times with PBS containing 1% Tween-20 (PBST) and incubated with fluorophore-conjugated secondary antibodies and 4’,6-diamidino-2-phenylindole (DAPI) for 1hr at 25°C. After staining, germ lines were washed three times with PBST. Slides were mounted by applying a drop of Fluoroshield mounting media (Sigma) on the germ lines followed by a coverslip. Stained germ lines were analyzed using the Zeiss Axiocam 40x objective or the Leica SP5 63x objective. Germ cell number and 3D modelling were performed using Imaris 9.5. The diameter of DAPI stained nuclei in the mitotic region was defined as 2.5μm and a three-dimensional model was created to enable counting of nuclei. To quantify the number of cells in active mitosis, extruded germ lines were stained with an anti-pH3 antibody and pH3-positive nuclei counted.

EdU labelling was performed as in previous studies ^25, 47^. NGM agar plates (-peptone, 60μg/ml carbenicillin) were seeded with concentrated *E. coli* MG1693 that was grown in *E. coli* M9 minimal media (3g/L KH_2_PO_4_, 6g/L Na_2_HPO_4_, 0.5g/L NaCl, 1g/L NH_4_Cl, 2mM MgSO_4_, 0.1mM CaCl_2_, 0.4% glucose, 1.25μg/ml thiamin) supplemented with 0.5μM thymidine and 20μM EdU for 24h. Auxin and ethanol were added to NGM agar accordingly. Following EdU feeding and germline isolation, fixation was performed as above. The Click reaction was performed according to the manufacturer instructions (Invitrogen™, C10337), and DNA was stained by DAPI. Stained germ lines were analyzed using the Nikon C1 Confocal Microscope 60x objective.

### Analysis of germline apoptosis

Synchronized one-day-old adults and OP50 bacteria were washed with M9 buffer, and a final concentration of 50μM SYTO 12 (Thermo Scientific™) added to the worm/bacteria mixture. After incubation at 25°C for 5h, worms were transferred to a fresh seeded plate for 1h to allow the stained bacteria to be purged from the gut to reduce background fluorescence. Worms were then mounted on agarose pads for the scoring of apoptosis nuclei which emit green fluorescence.

### Fluorescence microscopy of *C. elegans*

Animals were anesthetized with 20mM NaN_3_ on 5% agarose pads, and images were obtained with an Axio Imager M2 fluorescence microscope and Zen software (Zeiss).

### RNA Sequencing

On entering adulthood, a synchronized population of *mog-7::degron::GFP; sun-1p::TIR1* hermaphrodites were evenly divided into three groups (~1000 animals per sample), with two groups incubated either on 1mM auxin or ethanol (control) for 2h. After 1h, the remaining group was incubated on 1mM auxin for 1h. Six replicates of each sample were collected for RNA extraction and sequencing.

### Bioinformatics methods

Raw reads were first trimmed for adapter content and low-quality base-calls using Trimmomatic ^48^ in palindrome mode, using a sliding window of 5nt with quality threshold 15, removing any reads < 40nt after the trimming. Cutadapt was then used to remove reads with SL-1 and SL-2 spliced-leader sequences, followed by a first-pass mapping of reads to the WBCel235.99 genome using STAR ^49^, with conservative parameters for novel splice sites (alignSJDBoverhangMin 10). The splice junction files from this first-pass mapping were used with more liberal settings in STAR (alignSJDBoverhangMin 4) in a second-pass alignment of the reads. Duplicate alignments were marked with Picard MarkDuplicates and alignments sorted with Samtools ^50^.

These unfiltered alignments were used to analyze retained introns. A custom annotation was produced that only contained intronic sequences from the WBCel235.99 annotation as counting-bins, and featureCounts ^51^ was then used to count any read ovelapping an intron by more than 3 nt. The EdgeR quasi-likelihood method ^52^ was used to identify introns that were differentially represented in the 1h auxin-treated samples. Only introns with at least 5 reads in one of the samples and at least 3 samples with counts-per-million-intronic-reads above 0.5 were included for analysis. Because the auxin-treated samples were so skewed towards intronic sequences, and only intronic alignments were counted in each library, a custom library-size normalization was used whereby the log-fold-change (treated - untreated) was modeled as two normal distributions (using the function normalMixEM in the R package mixtools) and only the introns in the lower foldchange population (about 9000 introns) were used to generate library normalization factors. 17,647 introns in 5,014 genes were identified as differentially present in 1h treated samples, at a false discovery rate of 0.001. Apart from analysis of specific introns, the unfiltered alignments were also used to assess the genome-wide increase in non-exonic mRNA in the treated animals (Fig. 4 and Fig. S5).

Strictly exonic versions of the read alignments were generated in order to assess differences in gene expression and canonical splice site selection between auxin-treated and control samples, without interference from the significant amounts of non-canonical transcript / intronic / intergenic RNA found in the treated worms. Custom scripts were used to remove reads if an alignment overlaps any non-exonic region by more than 3 nt, according to the WBCel235.99 annotation. For calling differentially expressed genes, featureCounts ^51^ was used to count read alignments per-gene and Limma Voom ^53^ in Degust was used to perform the statistical analysis.

### RT-PCR assays

Total RNA was isolated using the RNeasy Mini Kit (QIAGEN 74104), according to manufacturer instructions. 500ng of RNA was reverse-transcribed to cDNA using 0.5mg/ml oligodT primers and the ImProm-II Reverse Transcription System (A3800) followed. cDNA was diluted to 1:5 in RNAse-free water. RT-PCR amplicons were run on a standard DNA electrophoresis gel.

### Quantification and statistical analysis

All experiments were performed in three independent replicates and the experimenter was blinded to genotype. Statistical analysis was performed in GraphPad Prism 7 using one-way analysis of variance (ANOVA) for comparison followed by Dunnett’s Multiple Comparison Test where applicable. Welch’s t-test was performed if the comparison was for two conditions. Values are expressed as mean ± s.e.m. Differences with a *P* value <0.05 were considered significant.

## ACKNOWLEDGEMENTS

We thank Judith Yanowitz, Peter Boag and members of the Pocock Laboratory for comments on the manuscript. We thank Hannah Seidel for advice and custom R script for germ cell analysis. Some strains were provided by the *Caenorhabditis* Genetics Center (University of Minnesota), which is funded by NIH Office of Research Infrastructure Programs (P40 OD010440).

## Author contributions

Conceptualization, W.C., S.G. and R.P.; Methodology, W.C., S.A., S.G. and R.P.; Investigation, W.C., C.T., S.A. and S.G.; Writing - Original Draft, R.P.; Writing - Review & Editing, W.C., C.T., S.A., S.G. and R.P.; Funding Acquisition, S.G., R.P.; Resources, S.G. and R.P; Supervision, S.G. and R.P.

## Data and materials availability

Data is available in the main text or the supplementary materials. Sequencing data is available under NCBI accession number GSE162055.

